# Targeting the Myeloid Immune Checkpoint ILT3 (LILRB4) with Small Molecules Enables Reprogramming of Suppressive Tumor Immunity

**DOI:** 10.64898/2026.06.05.730341

**Authors:** Somaya A. Abdel-Rahman, Antonio Monari, Tom Miclot, Florent Barbault, Moustafa T. Gabr

## Abstract

Cancer immunotherapy has transformed cancer treatment; however, durable responses remain limited by suppressive myeloid populations within the tumor microenvironment. Leukocyte immunoglobulin-like receptor B4 (LILRB4/ILT3) is an emerging myeloid immune checkpoint implicated in immune evasion and resistance to immunotherapy, yet small molecule targeting of ILT3 remains largely unexplored. Here, we report the discovery of small molecule ILT3 modulators identified using a Dianthus-based temperature-related intensity change (TRIC) screening platform. Screening of an 8,961-member Enamine Library identified multiple direct ILT3 binders, with lead compound **ICB-7** demonstrating high-affinity binding to recombinant human ILT3 by microscale thermophoresis and robust cellular target engagement in CETSA assays. Molecular docking and molecular dynamics simulations revealed a stable hydrophobic binding pocket within the D2 domain of ILT3. Functionally, **ICB-7** disrupted the ILT3-SCG2 interaction and inhibited downstream SHP1, SHP2, and STAT3 signaling. In patient-derived colorectal cancer and acute myeloid leukemia co-culture models, **ICB-7** enhanced cytotoxic T-cell activity, and reduced tumor-cell viability. The compound also demonstrated favorable pharmacokinetic and safety properties together with significant anti-tumor efficacy in the CT26 syngeneic colorectal carcinoma model. Collectively, these findings establish ILT3 as a tractable target for small-molecule immunomodulation and support pharmacological targeting of suppressive myeloid checkpoints as a promising cancer immunotherapy strategy.

## INTRODUCTION

Cancer immunotherapy has transformed the treatment landscape across multiple malignancies; however, a substantial proportion of patients fail to achieve durable responses due to the establishment of profoundly immunosuppressive tumor microenvironments.^1-4^ While current immune checkpoint therapies primarily target T cell-associated pathways such as PD-1/PD-L1 and CTLA-4, growing evidence indicates that suppressive myeloid populations including tumor-associated macrophages (TAMs), myeloid-derived suppressor cells (MDSCs), and tolerogenic dendritic cells play central roles in limiting anti-tumor immunity and driving resistance to immunotherapy.^5-9^ These populations promote immune evasion through inhibitory cytokine production, impaired antigen presentation, suppression of cytotoxic T-cell activation, and remodeling of the tumor microenvironment toward tolerogenic states.^10-12^ Consequently, pharmacological modulation of suppressive myeloid checkpoints has emerged as an important strategy for overcoming resistance to cancer immunotherapy and restoring productive anti-tumor immune responses.^13-15^

Among emerging myeloid checkpoints, leukocyte immunoglobulin-like receptor B4 (LILRB4, also known as ILT3) has gained significant attention as a potent regulator of immune suppression in both hematologic malignancies and solid tumors.^16-20^ ILT3 is highly expressed on suppressive myeloid populations including TAMs, MDSCs, dendritic cells, and monocytic leukemia cells, where it promotes tolerogenic phenotypes and suppresses T-cell activation.^21-24^ Upon activation, ILT3 signals through immunoreceptor tyrosine-based inhibitory motifs (ITIMs), leading to recruitment of SHP1 and SHP2 phosphatases and subsequent suppression of inflammatory and cytotoxic immune programs.^25-27^ Elevated ILT3 expression has been associated with poor prognosis, T-cell dysfunction, resistance to checkpoint blockade, and enhanced tumor progression across multiple cancer types.^28-30^ Preclinical studies using antibody-mediated inhibition of ILT3 have demonstrated restoration of T-cell activity, remodeling of suppressive myeloid phenotypes, and enhanced anti-tumor immunity, validating ILT3 as a promising immunotherapeutic target.^31-33^ Despite these advances, therapeutic efforts targeting ILT3 have remained largely restricted to biologics, while small-molecule modulation of ILT3 remains substantially underexplored.

Small molecules offer several potential advantages for targeting suppressive immune checkpoints, including improved tumor penetration, oral bioavailability, tunable pharmacokinetic properties, and the ability to access signaling mechanisms that may be less tractable using antibody-based approaches.^34-36^ Recent studies, including our own work,^37-45^ have demonstrated the feasibility of pharmacologically modulating immune checkpoint pathways using first-in-class small molecules targeting protein-protein interactions within the tumor immune microenvironment. However, discovery of small-molecule modulators targeting ILT3 remains challenging due to the non-enzymatic nature of the receptor and the absence of robust activity-based screening platforms suitable for direct target engagement studies. To address this challenge, we implemented a Dianthus-based temperature-related intensity change (TRIC) screening platform to identify compounds capable of directly engaging ILT3. Hits emerging from the primary screening cascade were subsequently progressed through orthogonal biophysical validation, computational analysis, and functional immune profiling.

Here, we report the identification of direct small molecule ILT3 modulators that reverse suppressive myeloid immune signaling and restore anti-tumor immune activity. Biophysical studies confirmed high-affinity target engagement, while functional assays demonstrated modulation of ILT3-associated signaling pathways and reprogramming of suppressive myeloid phenotypes. In human patient-derived immune cell systems, pharmacological targeting of ILT3 enhanced T-cell activation and promoted inflammatory immune responses. Furthermore, treatment with the lead compound significantly suppressed tumor growth and remodeled the tumor immune microenvironment in the CT26 syngeneic colon carcinoma model in vivo. Together, these findings establish ILT3 as a tractable target for small molecule immunomodulation and support pharmacological targeting of suppressive myeloid checkpoints as a promising strategy for cancer immunotherapy.

## RESULTS

### Dianthus Screening Identifies Direct Small-Molecule ILT3 Modulators

To identify small molecules capable of directly engaging ILT3, we implemented a Dianthus-based temperature-related intensity change (TRIC) screening workflow using recombinant human ILT3 extracellular domain (Figure 1A). Because ILT3 is a non-enzymatic immune receptor lacking conventional catalytic activity, direct target engagement approaches were prioritized over activity-based screening strategies. The Enamine Protein Mimetic Library, consisting of 8,961 structurally diverse compounds, was screened under solution-phase conditions at a final concentration of 10 µM in the presence of fluorescently labeled ILT3 protein. Binding-associated fluorescence perturbations were quantified as normalized fluorescence changes (ΔF_norm_) relative to vehicle-treated controls.

**Figure 1.**
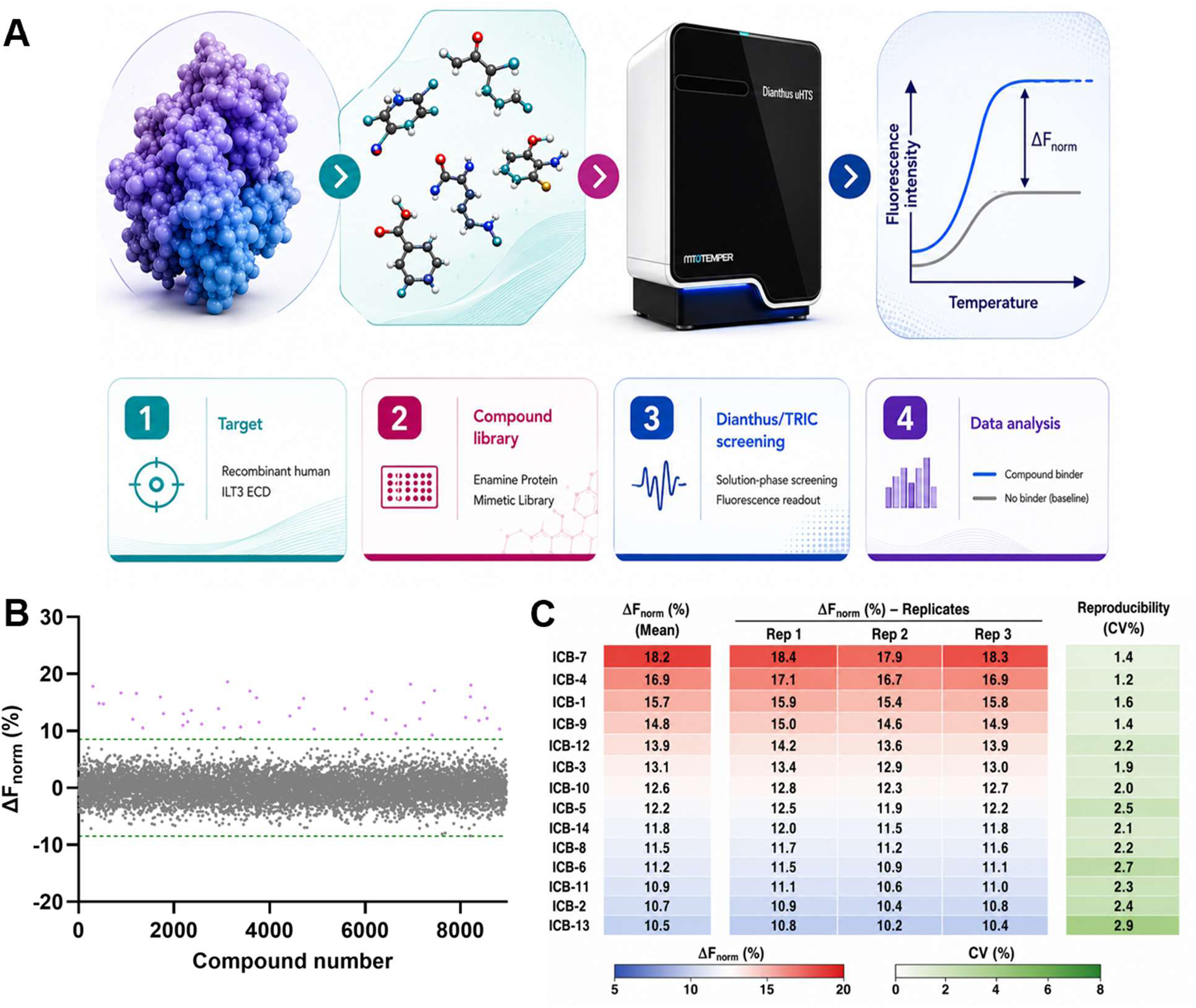
Dianthus/TRIC screening workflow and secondary validation of candidate ILT3 binders. **(A)** Schematic overview of the Dianthus-based temperature-related intensity change (TRIC) screening workflow used to identify direct small molecule binders of ILT3. Recombinant human ILT3 extracellular domain was screened against the Enamine Protein Mimetic Library under solution-phase conditions using fluorescently labeled protein. Binding-induced fluorescence perturbations were quantified as normalized fluorescence changes (ΔF_norm_), followed by hit identification, refinement, and progression into orthogonal validation studies. **(B)** Primary Dianthus/TRIC screening results for the 8,961-member Enamine Protein Mimetic Library. Compounds exceeding the predefined hit threshold of 3 standard deviations above vehicle-treated controls are highlighted in purple. **(C)** Heatmap summarizing secondary validation of 14 candidate ILT3 binders advanced from the primary screen. Mean ΔF_norm_ values, technical replicates, and coefficients of variation (CV%) are shown to assess assay reproducibility and signal consistency across validated candidates.

Primary screening identified 46 preliminary hits exceeding the predefined hit threshold of 3 standard deviations above vehicle-treated controls (Figure 1B). Following removal of compounds displaying signal instability, aggregation behavior, or poor reproducibility, 14 structurally distinct candidates were progressed into secondary validation studies. Heatmap analysis of the advanced candidates demonstrated strong assay reproducibility and consistent ΔF_norm_ responses across technical replicates (Figure 1C). Several compounds exhibited reproducible interaction profiles consistent with direct ILT3 engagement and were subsequently progressed into orthogonal biophysical characterization.

### Orthogonal Biophysical Studies Validate Direct ILT3 Engagement

To quantitatively evaluate direct interactions with ILT3, the top 8 compounds emerging from the Dianthus/TRIC screening workflow were further analyzed by microscale thermophoresis (MST) using recombinant human ILT3 extracellular domain protein. Four compounds (Table S1) produced reproducible, concentration-dependent binding signals indicative of direct target association. Among these, **ICB-7** (Figure 2A) and **ICB-9** (Figure 2B) displayed the highest binding affinities, with equilibrium dissociation constants (K_D_) values of 156 ± 11.8 nM (Figure 2C) and 887 ± 36.1 nM (Figure 2D). The MST traces demonstrated well-defined saturation at elevated ligand concentrations together with stable fluorescence behavior across replicates, consistent with specific ILT3 engagement rather than nonspecific aggregation effects.

**Figure 2.**
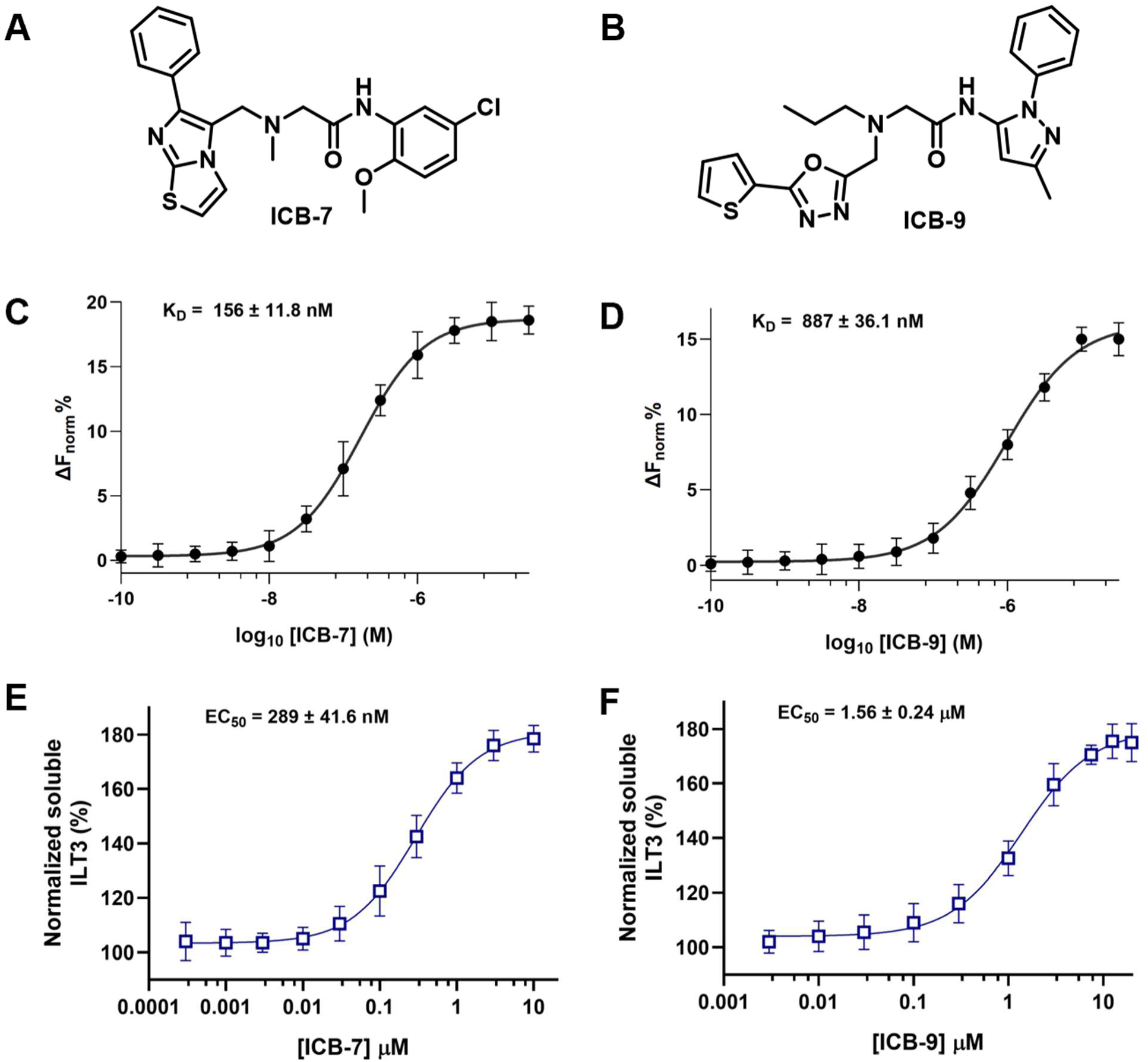
Identification and validation of direct ILT3/LILRB4-binding small molecules. **(A,B)** Chemical structures of the validated hit compounds **ICB-7** and **ICB-9** identified from the screening campaign. **(C,D)** MST analysis demonstrating direct binding of **ICB-7** and **ICB-9** to recombinant human ILT3/LILRB4 protein. **ICB-7** exhibited a K_D_ value of 156 ± 11.8 nM, whereas **ICB-9** showed a weaker binding affinity with a K_D_ value of 887 ± 36.1 nM. Data are presented as ΔF_norm_ (%) values plotted against compound concentration. **(E,F)** CETSA analysis demonstrating dose-dependent stabilization of endogenous ILT3/LILRB4 in cells treated with **ICB-7** or **ICB-9**. **ICB-7** induced stronger cellular target engagement with an EC_50_ value of 289 ± 41.6 nM, while **ICB-9** exhibited an EC_50_ value of 1.56 ± 0.24 μM. Data are presented as normalized soluble ILT3 levels following thermal challenge. Values represent mean ± SD (n=5).

To assess whether **ICB-7** and **ICB-9** also interact with ILT3 in a cellular setting, a dose-dependent cellular thermal shift assay (CETSA) was performed using our optimized ILT3 CETSA platform. ILT3-expressing cells were incubated with increasing concentrations of the tested compound for 60 min prior to thermal challenge at 51 °C for 3 min, followed by quantification of soluble ILT3 protein. Treatment with **ICB-7** and **ICB-9** produced concentration-dependent thermal stabilization of ILT3, with an EC_50_ values of 289 ± 41.6 nM and 1.56 ± 0.24 μM for **ICB-7** and **ICB-9**, respectively (Figure 2E and 2F), supporting direct intracellular target engagement and demonstrating that the compound remains active under native cellular conditions. Together, these results highlight the effectiveness of the Dianthus/TRIC discovery platform for the identification and validation of direct small molecule binders targeting ILT3.

### Computational Characterization of ILT3 Ligand Binding

To gain mechanistic insight into ligand recognition by ILT3, molecular docking and molecular dynamics (MD) simulations were performed to characterize the binding modes and stability of **ICB-7** and **ICB-9**. Although the structure of ILT3 is available, no experimental structure of this protein in complex with an organic ligand has been reported to date. The crystallographic structure explicitly reveals the presence of a D1 domain spanning residues 1 to 96 and a D2 domain extending from residues 97 to 196. The overall structure is predominantly composed of β-sheets connected through an interdomain interface characterized by an angle of approximately 107°.

In order to identify potential interaction modes for **ICB-7** and **ICB-9**, molecular docking calculations were performed over the entire protein surface, leading to the generation of 2000 docking poses. Remarkably, despite the exhaustive exploration of the ILT3 surface, analysis of the 40 best-ranked poses obtained for both ligands revealed only two preferential binding regions. Figure 3 illustrates these two binding pockets located within the D1 and D2 domains, while Table 1 summarizes the distribution of docking poses associated with each binding site.

**Figure 3.**
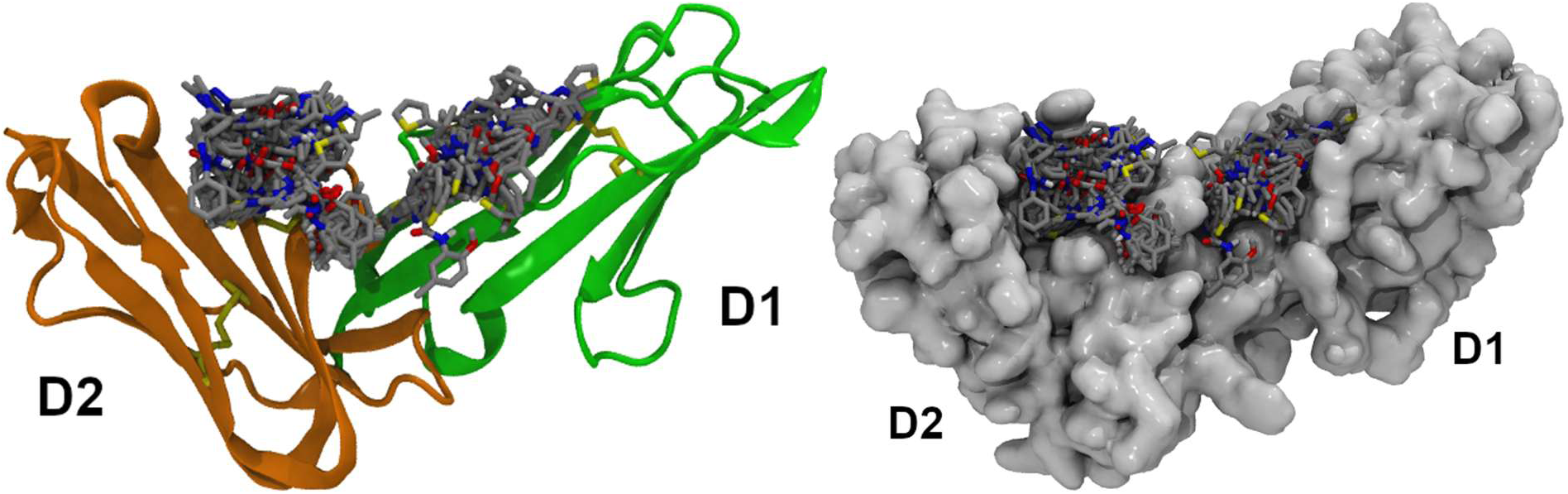
Illustration of the two biding modes observed for the ILT3 structures for ILT3 for **ICB-7** and **ICB-9** with ribbons representation (left) or solvent accessible surface (right). D1 and D2 refers to the two domains of ILT3 structure.

**Table 1.**
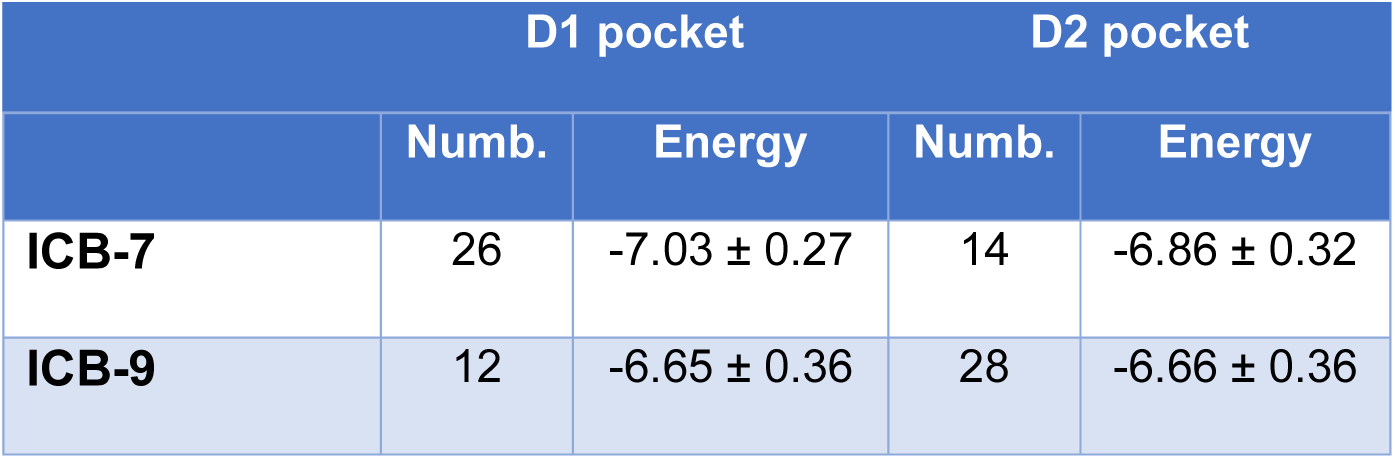
Molecular docking results for **ICB-7** and **ICB-9**. Numb. indicates the numbers of docking poses which fall into D1 or D2 pocket. Energies are in kcal/mol.

The molecular docking results therefore clearly identify only two possible interaction modes, hereafter referred to as the D1 pocket and D2 pocket, according to their localization within the D1 and D2 domains, respectively. Examination of the number of docking poses obtained for both compounds indicates, as summarized in Table 1, that **ICB-7** exhibits a preference for the D1 pocket, whereas **ICB-9** preferentially populates the D2 pocket. Nevertheless, no strict exclusivity between the two binding regions is observed.

Similarly, analysis of the average docking energies associated with each pocket for both compounds reveal very similar values which, considering the corresponding standard deviations, cannot be discriminated on the basis of energetic criteria alone. Consequently, in order to further evaluate the relevance and stability of these predicted binding modes, the poses exhibiting the best interaction energies for each pocket (D1 and D2) were selected for MD simulations. In contrast to docking calculations, which provide a largely static description of ligand recognition, MD simulations make it possible to account for the intrinsic flexibility of both the ligand and the protein, the influence of solvent effects, as well as the temporal stability of intermolecular interactions. This approach therefore enables a more realistic evaluation of the persistence and structural consistency of the predicted ligand/protein complexes.

Accordingly, four independent 500 ns MD simulations were performed under NPT conditions by selecting the best-ranked docking poses obtained for each compound within the D1 and D2 pockets. Monitoring of the potential, kinetic, and total energies, together with the simulation box volumes, confirmed the overall stability of all systems throughout the simulations. Subsequent analyses were carried out over the last 100 ns of each trajectory. The behavior of **ICB-7** and **ICB-9** during the simulations is of particular interest, and Figure 4 presents the evolution of the ligand root mean square deviations (RMSD) values throughout the MD trajectories.

**Figure 4.**
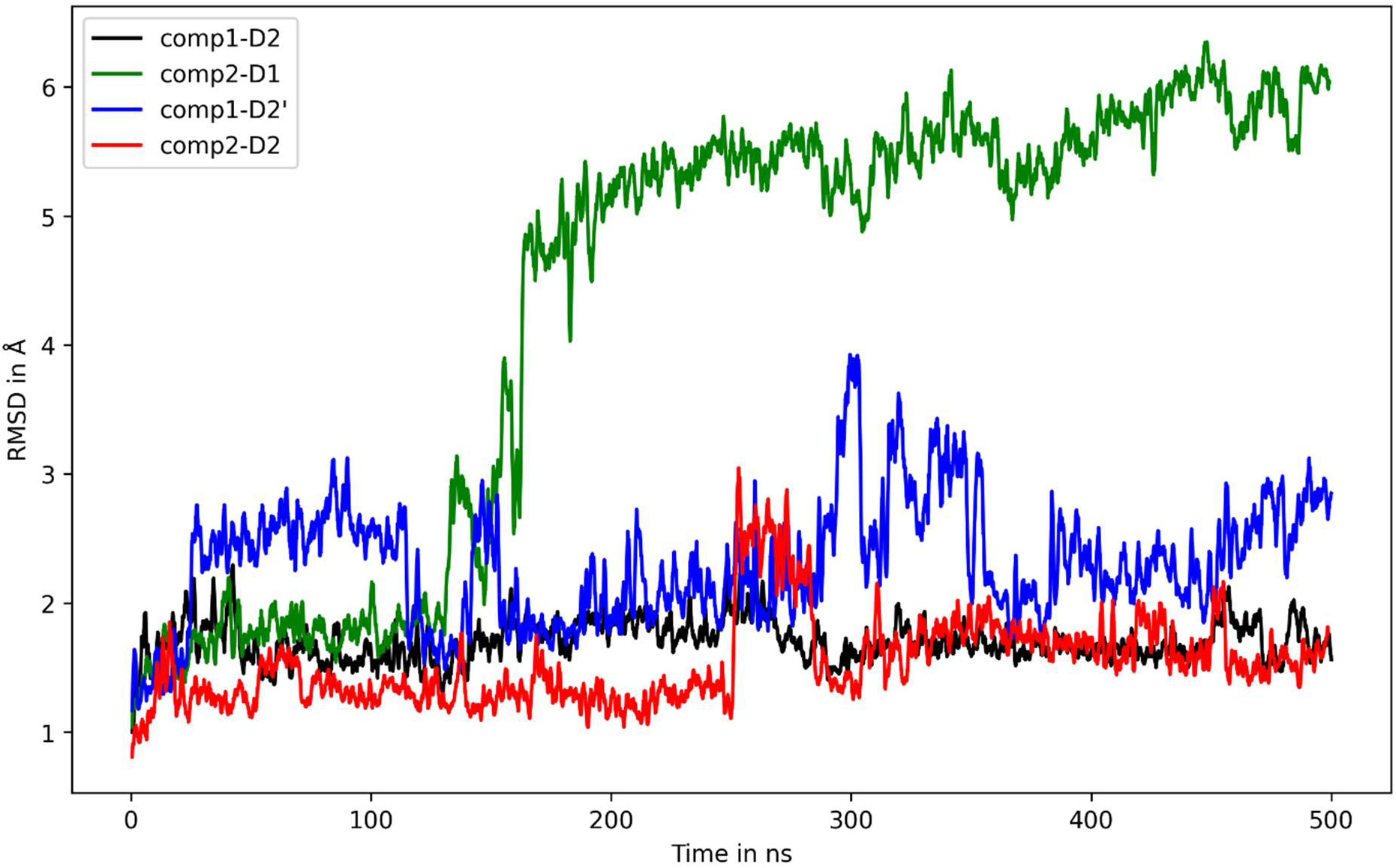
RMSD variations versus trajectory time for the four simulations for the interactions between **ICB-7** (comp1) and **ICB-9** (comp2) with ILT3.

For **ICB-7**, an interesting behavior was observed during the simulations. The ligand initially positioned within the D1 pocket was not stable over time and quickly dissociated from this binding site after approximately 25 ns. The compound subsequently migrated toward another interaction region located on the D2 domain, which does not exactly correspond to the previously identified D2 pocket. The ligand remained stably associated with this new interaction region, hereafter referred to as D2’, for the remainder of the simulation. This behavior is testified by observing its RMSD curve (blue line in Figure 4) where an increase is observed of **ICB-7** after 25 ns. In contrast, the **ICB-7** ligand initially positioned within the D2 pocket remained stable in this region and exhibited only limited structural fluctuations throughout the MD trajectory. Again, this behavior can also being observed on the RMSD curve (black line in Figure 4) where it can be seen that the **ICB-7** fluctuation along the trajectory time is moderate and never up to 2 Å which suggests a particular stable interaction onto the D2 domain of ILT3.

A particularly interesting behavior was observed for **ICB-9**. Indeed, the ligand initially positioned within the D1 pocket was not stable and dissociated from its interaction site after approximately 130 ns of simulation time. This behavior is therefore similar to that observed for **ICB-7** within the same interaction region. However, unlike **ICB-7**, **ICB-9** was unable to identify an alternative stable interaction site and progressively detached from ILT3, as evidenced by the rapid increase of its RMSD after 130 ns of trajectory (green curve in Figure 4). Following this dissociation event, a continuous increase in the RMSD was observed, indicating that the ligand diffuses along the surface of ILT3 without establishing a stable interaction with the protein during the remaining simulation time. In contrast, the behavior of **ICB-9** within the D2 pocket was markedly different, as the compound remained stably bound to this interaction site throughout the simulation. Similarly to **ICB-7**, this stability is reflected by the RMSD evolution along the trajectory, which consistently remained below 2 Å (red curve in Figure 4).

To quantify the strength of the interactions between the ligands and ILT3, MM/GBSA calculations were subsequently performed. The corresponding binding free energy values are summarized in Table 2. This calculation was not performed for **ICB-9** in the D1 pocket since the ligand dissociated from the ILT3 protein during the MD simulation.

**Table 2.** Free energy computations made for **ICB-7** in the D2’ and D2 pocket and **ICB-9** n the D2 pocket. All energies are in kcal/mol.

| Component | ICB-7-D2' | ICB-7-D2 | ICB-9-D2 |
| --- | --- | --- | --- |
| **DE<sub>vdw</sub>** | -24.67 ± 0.18 | -47.43 ± 0.11 | -33.18 ± 0.24 |
| **DE<sub>elec</sub>** | -08.41 ± 0.47 | -13.90 ± 0.15 | -07.37 ± 0.33 |
| **DG<sub>solv</sub>** | +19.54 ± 0.37 | +27.86 ± 0.14 | +17.99 ± 0.30 |
| **DG<sub>tot</sub>** | -13.54 ± 0.13 | -33.43 ± 0.12 | -22.56 ± 0.24 |

On Table 2, comparison of the interaction free energies calculated for **ICB-7** reveals a substantial difference between the D2 pocket and the D2’ interaction site, with binding free energies of -33.4 and -13.5 kcal/mol, respectively. Considering that the D2’ site was not identified during the docking calculations and only emerged from ligand migration during the molecular dynamics simulation, these results strongly suggest that D2’ does not correspond to the preferential binding site of **ICB-7**. Consequently, the present simulations support the existence of a single preferential interaction site for both **ICB-7** and **ICB-9**, namely the D2 pocket. The calculated binding free energies for this interaction site are strongly favorable, reaching -33.43 and -22.56 kcal/mol for c **ICB-7** and **ICB-9**, respectively.

Analysis of the different energetic contributions indicates that ligand binding results from a combination of a limited electrostatic contribution together with a dominant van der Waals component, partially counterbalanced by a positive solvation free energy term. Such an energetic profile is generally characteristic of ligand binding within a predominantly hydrophobic cavity. Representative structures obtained from the MD simulations of **ICB-7** and **ICB-9** in complex with ILT3 are presented in Figure 5.

**Figure 5.**
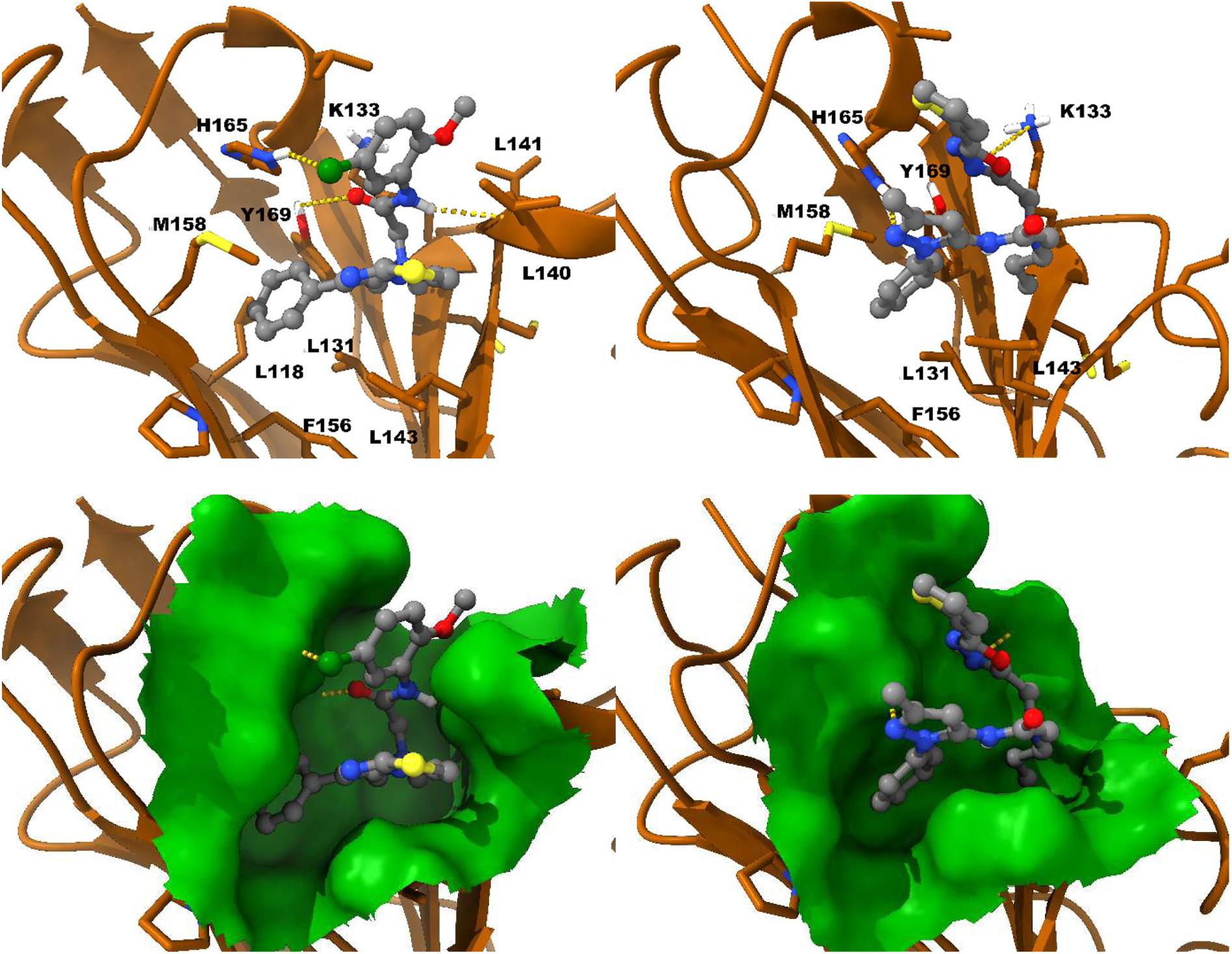
Representative structures illustrating the interactions between the ILT3 protein and the ligands **ICB-7** (left panel) and **ICB-9** (right panel). The ligands are displayed using a ball-and-stick representation with carbon atoms shown in gray, whereas the protein residues involved in the interactions are represented as sticks with orange carbon atoms. Yellow dashed lines indicate favorable interactions established between the ligands and ILT3. The lower panel presents the molecular interaction surface, revealing the presence of a cavity within the protein into which the ligands are inserted.

For **ICB-7**, three hydrogen bonds were identified involving the imidazole side chain of H165 and the chlorine atom of the aromatic ring of **ICB-7**, the phenolic group of Y169 and the carbonyl oxygen of the amide function of **ICB-7**, as well as the amide NH group of **ICB-7** interacting with the backbone oxygen atom of L140. These three hydrogen bonds strongly anchor the ligand within the binding site, thereby promoting the insertion of the phenyl moiety of **ICB-7** into a hydrophobic cavity. As illustrated in the lower-left panel of Figure 5, this cavity is mainly formed by the residues L118, L131, F156, and M158.

For **ICB-9**, only two interactions, explaining this way a lowest free energy of association in table T2, were identified as responsible for stabilizing the ligand within the same hydrophobic cavity. Nevertheless, as shown in the upper-right panel of Figure 5, it is noteworthy that the same residues are involved, highlighting the strong affinity of this region of ILT3 toward ligand binding. Similarly to **ICB-7**, a hydrogen bond involving the imidazole side chain of H165 is observed with the pyrazole ring of **ICB-9**. In addition, a cation–π interaction is detected between the ammonium group of K133 and the 1,3,4-oxadiazole ring of **ICB-9**. This interaction is particularly interesting since similar cation–π interactions have previously been reported in anticancer compounds, notably within the binding site of combretastatin A4 on tubulin.^46^ These favorable interactions allow, as observed for **ICB-7**, the insertion of the phenyl ring of **ICB-9** into the same hydrophobic cavity of ILT3. The fact that both ligands converge toward highly similar interaction modes, despite the absence of any prior constraints during the simulations, strongly supports the robustness and consistency of the proposed binding model and therefore increases the confidence in our computational results.

### ICB-7 Functionally Blocks the ILT3-SCG2 Signaling Axis and Attenuates Immunosuppressive Pathways

To investigate whether direct binding of **ICB-7** (the most potent compound from this study) to ILT3 translates into functional inhibition of receptor signaling, we examined its ability to interfere with the interaction between ILT3 and secretogranin II (SCG2), a ligand associated with suppressive myeloid signaling and tumor immune escape. In a TR-FRET competition assay, **ICB-7** inhibited the ILT3-SCG2 interaction in a concentration-dependent fashion, yielding an IC_50_ value of 694 ± 32.4 nM (Figure 6A), consistent with effective disruption of this inhibitory ligand-receptor interaction.

**Figure 6.**
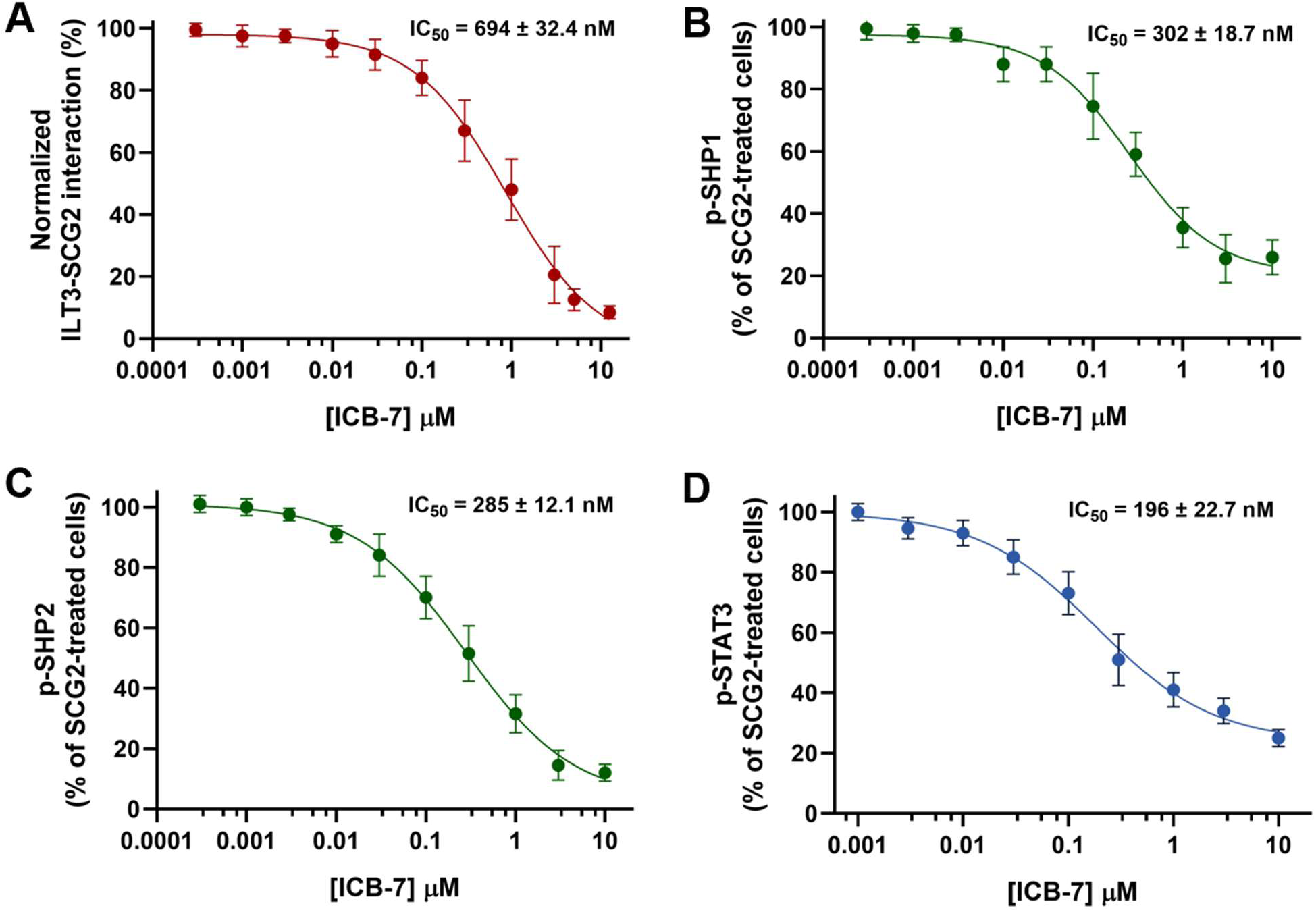
ICB-7 disrupts ILT3-SCG2 engagement and inhibits downstream suppressive signaling. **(A)** TR-FRET competition assay demonstrating concentration-dependent inhibition of the ILT3-SCG2 interaction by **ICB-7**. **(B-D)** ELISA-based analysis of intracellular signaling pathways in ILT3-expressing THP-1-derived macrophage-like cells stimulated with SCG2 and treated with increasing concentrations of **ICB-7**. The compound produced dose-dependent inhibition of **(B)** phospho-SHP1, **(C)** phospho-SHP2, and **(D)** phospho-STAT3 relative to SCG2-stimulated controls. Collectively, these findings demonstrate that **ICB-7** suppresses ligand-dependent ILT3 signaling and downstream immunoregulatory pathways. Data are shown as mean ± SD (n = 5). IC_50_ values were determined by nonlinear regression analysis.

Because ILT3 transduces inhibitory signals through recruitment of SHP-family phosphatases via intracellular ITIM motifs, we next assessed whether **ICB-7** modulates early downstream signaling events triggered by SCG2. In ILT3-expressing THP-1-derived macrophage-like cells, SCG2 treatment induced robust phosphorylation of SHP1 and SHP2, confirming activation of canonical inhibitory signaling pathways. **ICB-7** significantly reduced SCG2-driven SHP1 and SHP2 phosphorylation in a dose-dependent manner, with IC_50_ values of 302 ± 18.7 nM and 285 ± 12.1 nM, respectively (Figure 6B,C), indicating effective suppression of proximal ILT3 signaling.

We subsequently examined the impact of **ICB-7** on STAT3 activation, a central mediator of suppressive myeloid phenotypes and tumor-associated immune dysfunction. SCG2 stimulation markedly increased STAT3 phosphorylation in ILT3-positive cells, whereas treatment with **ICB-7** strongly decreased p-STAT3 levels in a concentration-dependent manner (IC_50_ = 196 ± 22.7 nM, Figure 6D). Together, these data demonstrate that **ICB-7** inhibits ILT3-mediated suppressive signaling at multiple mechanistic levels, including ligand binding, SHP phosphatase activation, and downstream STAT3 signaling. These findings support the potential of **ICB-7** as a functional small molecule antagonist capable of counteracting ILT3-dependent immunosuppressive pathways involved in tumor immune evasion.

### ICB-7 Restores Anti-Tumor Immune Activity in Patient-Derived Solid and Hematologic Tumor Co-Culture Models

To determine whether inhibition of ILT3 by **ICB-7** can restore immune function in clinically relevant systems, we evaluated the compound in ex vivo co-culture models representing both solid tumors and hematologic malignancies. In colorectal cancer models, PBMCs isolated from colorectal cancer patients were co-cultured with HCT116 or HT-29 tumor cells under SCG2 stimulation to induce suppressive ILT3 signaling. SCG2 exposure significantly impaired anti-tumor immune responses, as reflected by reduced IFN-γ and IL-2 secretion and increased tumor cell survival (Figure 7A-C for HCT116; Figure S1A-C for HT-29). Treatment with **ICB-7** restored cytokine secretion in a dose-dependent manner (1-25 μM) and reduced tumor viability, consistent with recovery of immune-mediated tumor killing. At higher concentrations, the activity of **ICB-7** approached the effects observed with a blocking anti-ILT3 antibody used as a target-specific control.

**Figure 7.**
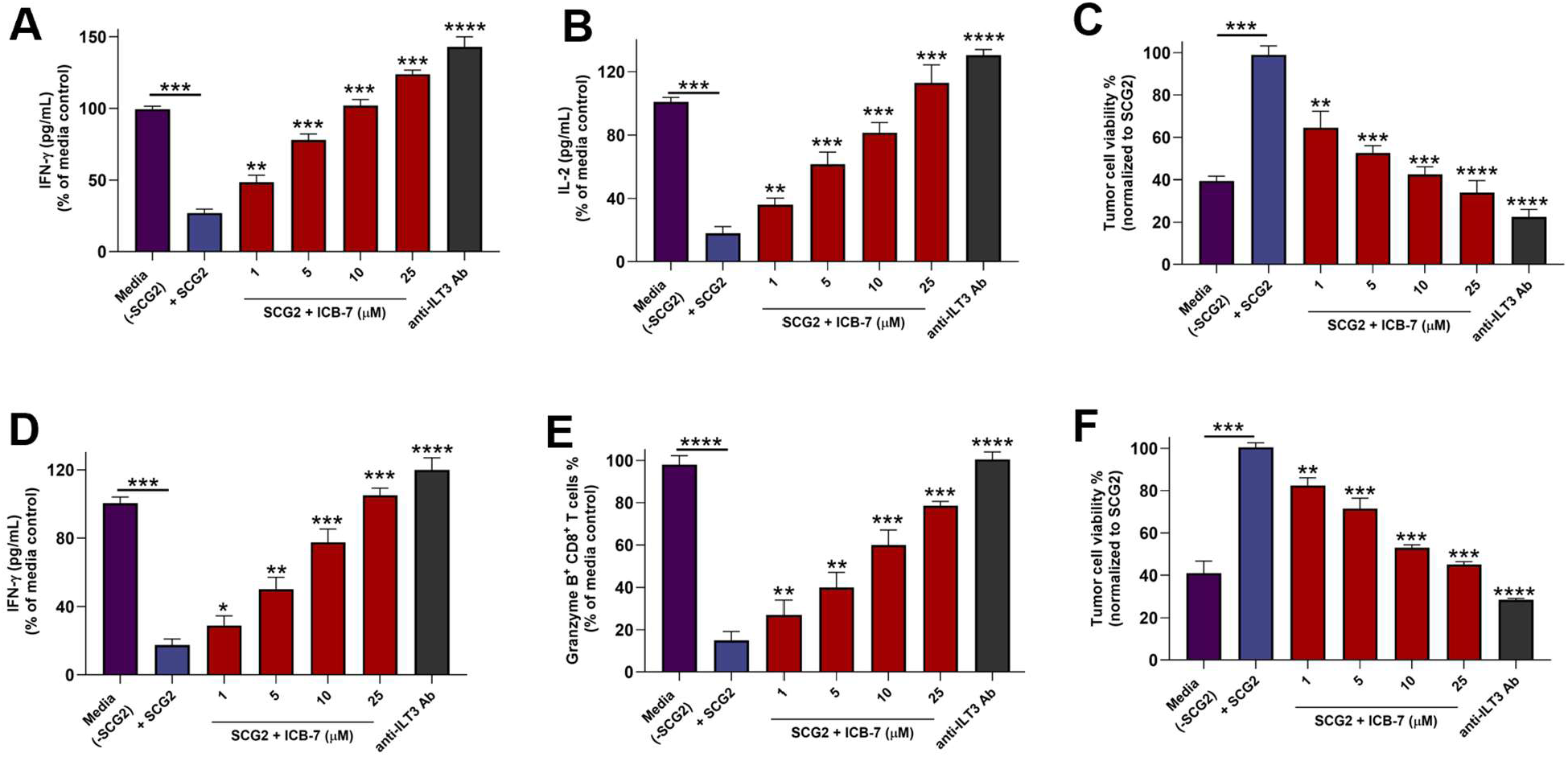
ICB-7 reverses SCG2-mediated immunosuppression in colorectal cancer and AML co-culture models. (A,B) In HCT116 colorectal cancer co-cultures, SCG2 suppressed anti-tumor immune responses, as demonstrated by decreased IFN-γ and IL-2 secretion, whereas **ICB-7** restored cytokine production in a dose-dependent manner. **(C) ICB-7** also reduced HCT116 tumor-cell viability in SCG2-treated co-cultures, consistent with enhanced immune-mediated tumor killing. **(D)** In primary AML blast co-cultures, **ICB-7** restored IFN-γ secretion suppressed by SCG2-dependent ILT3 signaling. **(E)** Treatment with **ICB-7** increased the proportion of Granzyme B^+^ CD8^+^ T cells, indicating recovery of cytotoxic T-cell activity. **(F) ICB-7** significantly decreased the viability of primary AML blasts in SCG2-stimulated co-cultures. Across multiple functional assays, the activity profile of **ICB-7** approached that observed with a blocking anti-ILT3 antibody used as a target-specific positive control. Data are presented as mean ± SD from independent donors analyzed in technical replicates (n=5). Statistical significance was evaluated using one-way ANOVA followed by Tukey’s multiple-comparisons test. \*\**p* < 0.05, \*\*\**p* < 0.001, \*\*\*\**p* < 0.0001 versus media + SCG2 controls.

To determine whether these effects extend beyond solid tumors, **ICB-7** was further evaluated in AML co-culture systems using THP-1 cells or primary AML blasts cultured with PBMCs or enriched CD3^+^ T cells. Similar to the colorectal cancer setting, SCG2 induced a suppressive phenotype characterized by decreased IFN-γ production, impaired cytotoxic T-cell activation, and enhanced AML cell survival (Figure 7D-F for AML blasts; Figure S2D-F for THP-1 cells). **ICB-7** effectively reversed these suppressive effects in a concentration-dependent manner (1-25 μM), resulting in increased IFN-γ secretion, elevated frequencies of Granzyme B^+^ CD8^+^ T cells, and reduced AML viability (Figure 7D-F; Figure S1D-F).

Importantly, **ICB-7** consistently recapitulated the activity profile of a blocking anti-ILT3 antibody across multiple orthogonal assays, strongly supporting an on-target mechanism of action. Collectively, these results demonstrate that **ICB-7** functions as a potent small molecule inhibitor of ILT3 capable of reversing SCG2-driven immunosuppression and restoring anti-tumor immune responses across diverse human tumor models.

### Pharmacokinetic (PK) Profiling of ICB-7

The preliminary PK and developability profiling of **ICB-7** demonstrated a favorable balance between physicochemical properties, metabolic stability, and safety-related parameters, supporting its suitability as a small molecule ILT3 modulator for further development (Table 3). **ICB-7** exhibited moderate lipophilicity with a LogD_7.4_ value of 3.24, consistent with compounds possessing balanced membrane permeability while avoiding excessive hydrophobicity that may compromise solubility or increase nonspecific binding. Despite its aromatic-rich scaffold, the compound maintained acceptable aqueous solubility, displaying a kinetic solubility of 48.6 μM in 1% DMSO/PBS and a FaSSIF solubility of 39.2 μM, indicating favorable solubilization under physiologically relevant intestinal conditions (Table 3).

**Table 3.** In vitro PK profile of ICB-7.

| PK parameter | ICB-7 |
| --- | --- |
| LogD<sub>7.4</sub><sup>a</sup> | 3.24 |
| Kinetic Solubility<br>1% DMSO/PBS<br>[μM]<sup>b</sup> | 48.6 |
| Solubility in<br>FaSSIF [μM] | 39.2 |
| PAMPA $P_{app}$<br>[cm/s] | $3.1 \times 10^{-6}$ |
| Stability in<br>simulated gastric<br>fluid (pH 1.2, 2h) | 94.3% |
| Stability in<br>simulated<br>intestinal fluid (pH<br>6.8, 2h) | 90.6% |
| $t_{1/2}$ [min] mouse<br>liver microsomes<sup>c</sup><br>/ CL<sub>int</sub><sup>d</sup> | 82.7 ± 6.2 /<br>21.8 ± 1.7 |
| $t_{1/2}$ [min] human<br>liver microsomes<sup>c</sup><br>/ CL<sub>int</sub><sup>d</sup> | 108.4 ± 7.5 /<br>15.6 ± 1.3 |
| $t_{1/2}$ [min] mouse plasma | >180 |
| $t_{1/2}$ [min] human plasma | >180 |
| $t_{1/2}$ [min] PBS | >300 |
| PPB human [%]<sup>c</sup> | 95.2 ± 0.64 |
| IC<sub>50</sub> against WI-38 [μM] | >100 |
| IC<sub>50</sub> against HS 27 [μM] | >100 |
| IC<sub>50</sub> against hERG [μM] | >100 |
| CYP3A4 inhibition (% at 10 mM) | 8.4 |
| CYP2D6 inhibition (% at 10 mM) | 4.1 |
| CYP2C9 inhibition (% at 10 mM) | 6.3 |
| CYP2C19 inhibition (% at 10 mM) | 5.0 |

<sup>a</sup>LogD (pH = 7.4) the distribution coefficient at physiological pH 7.4. <sup>b</sup>Kinetic solubility measured in 1% DMSO/PBS (pH 7.4) by UV-vis spectrophotometry (n=3). <sup>c</sup>Values are mean ± SD (n=3). <sup>d</sup>Intrinsic clearance expressed in μL/(min × mg) protein. Values are mean ± SD (n=3).

Permeability assessment using the PAMPA assay revealed a passive permeability coefficient (P_app_) of 3.1 × 10^−6^ cm/s, supporting the ability of **ICB-7** to traverse biological membranes efficiently. In parallel, the compound demonstrated strong chemical stability, remaining 94.3% intact following incubation in simulated gastric fluid (pH 1.2) for 2 h and 90.6% stable in simulated intestinal fluid (pH 6.8), suggesting resistance to degradation under gastrointestinal conditions. This profile may support future oral administration strategies.

Metabolic stability studies further highlighted the favorable microsomal stability of **ICB-7**. In mouse liver microsomes, the compound exhibited a half-life of 82.7 ± 6.2 min with an intrinsic clearance (CL_int_) of 21.8 ± 1.7 μL/(min × mg). Even greater stability was observed in human liver microsomes, where **ICB-7** demonstrated a half-life of 108.4 ± 7.5 min and a lower intrinsic clearance value of 15.6 ± 1.3 μL/(min × mg), supporting relatively slow metabolic turnover. Importantly, **ICB-7** also showed excellent plasma stability, remaining stable for more than 180 min in both mouse and human plasma and exceeding 300 min in PBS, indicating limited nonspecific degradation or chemical instability (Table 3).

Protein binding analysis revealed high plasma protein binding in human plasma (95.2 ± 0.64%), which is commonly observed for moderately lipophilic heteroaromatic compounds and may contribute to prolonged systemic exposure in vivo. Importantly, early safety profiling demonstrated an encouraging profile for **ICB-7**. The compound showed no measurable cytotoxicity against WI-38 lung fibroblasts or HS27 stromal cells at concentrations up to 100 μM. In addition, no significant hERG channel inhibition was detected at concentrations up to 100 μM, suggesting low predicted cardiotoxicity risk. CYP inhibition profiling further demonstrated minimal inhibition across major drug-metabolizing isoforms, including CYP3A4 (8.4%), CYP2D6 (4.1%), CYP2C9 (6.3%), and CYP2C19 (5.0%) at 10 μM, supporting a relatively low likelihood of CYP-mediated drug-drug interactions.

Collectively, these findings indicate that **ICB-7** combines favorable permeability, metabolic stability, chemical stability, and early safety characteristics with potent ILT3 target engagement, supporting its continued optimization and advancement as a promising immunomodulatory small-molecule scaffold.

To evaluate whether **ICB-7** achieves systemic exposure levels compatible with in vivo efficacy studies, PK profiling was performed in BALB/c mice following single-dose intravenous (IV) and oral administration. After IV dosing at 2 mg/kg, **ICB-7** displayed a plasma half-life of 4.6 h together with a systemic clearance of 9.4 mL/min/kg and a volume of distribution of 3.2 L/kg, indicating relatively slow clearance and broad tissue distribution. The corresponding plasma exposure (AUC0−∞) reached 10.9 µM·h, consistent with the favorable metabolic stability observed in liver microsomal assays.

Following oral administration at 25 mg/kg, **ICB-7** demonstrated efficient systemic absorption, reaching a plasma C_max_ of 3.8 µM with a T_max_ of 1.5 h. Oral dosing produced an AUC0−∞ of 52.7 µM·h, corresponding to an estimated oral bioavailability of approximately 48%. The terminal plasma half-life following oral administration was 5.1 h, supporting sustained systemic exposure throughout the dosing interval. Notably, plasma concentrations following oral dosing remained above the biochemical and cellular potency ranges determined in MST, CETSA, and TR-FRET signaling assays for several hours after administration, supporting effective systemic target engagement in vivo. Collectively, the favorable oral exposure profile, moderate plasma half-life, acceptable bioavailability, and clean preliminary safety characteristics of **ICB-7** supported the selection of once-daily oral dosing for subsequent efficacy studies in syngeneic tumor models.

### ICB-7 exhibits cross-species ILT3 engagement and suppresses colorectal tumor growth in vivo

Prior to evaluating the anti-tumor efficacy of **ICB-7** in vivo, we first determined whether the compound retained binding activity toward murine ILT3 to support pharmacological studies in syngeneic mouse tumor models. MST analysis confirmed direct interaction between **ICB-7** and recombinant murine ILT3, yielding a K_D_ of 262 ± 34.7 nM, demonstrating that the compound maintains high-affinity target engagement across species and is suitable for murine efficacy studies (Figure S2). Additional selectivity assessment using Dianthus/TRIC profiling revealed a strong and preferential fluorescence response for ILT3 relative to a panel of 12 related LILR family members and immune checkpoint proteins, supporting the selectivity of **ICB-7** toward its intended target (Figure S3).

We next evaluated the therapeutic potential of **ICB-7** in the CT26 syngeneic colorectal carcinoma model using immunocompetent BALB/c mice. CT26 tumor cells were implanted subcutaneously, and treatment was initiated once tumors reached approximately 75-100 mm^3^. Animals were randomized to receive either vehicle or **ICB-7** at 25 mg/kg by oral gavage once daily for 21 days. Dose selection was guided by the favorable PK properties of **ICB-7**, including sustained systemic exposure above the biochemical and cellular target-engagement concentrations associated with inhibition of ILT3 signaling pathways.

Oral administration of **ICB-7** produced significant inhibition of CT26 tumor progression compared with vehicle-treated controls, with clear divergence of tumor growth kinetics observed throughout the treatment period (Figure 8A). Consistent with these findings, final tumor weights collected at study termination were substantially reduced in the **ICB-7** treatment group relative to controls (Figure 8B). Importantly, treatment was well tolerated over the course of the study, with no significant changes in body weight or visible signs of systemic toxicity detected, in agreement with the favorable in vitro safety and ADME characteristics observed during early profiling studies. To determine whether tumor suppression by **ICB-7** was associated with restoration of anti-tumor immune activity in vivo, tumor lysates were analyzed for pharmacodynamic immune markers. Treatment with **ICB-7** significantly increased intratumoral IFN-γ and IL-2 levels compared with vehicle-treated tumors, indicating enhanced immune activation within the tumor microenvironment (Figure 8C,D). These in vivo findings are consistent with the ex vivo co-culture studies and support a mechanism in which pharmacological inhibition of ILT3 alleviates suppressive myeloid signaling, thereby restoring productive anti-tumor immune responses.

**Figure 8.**
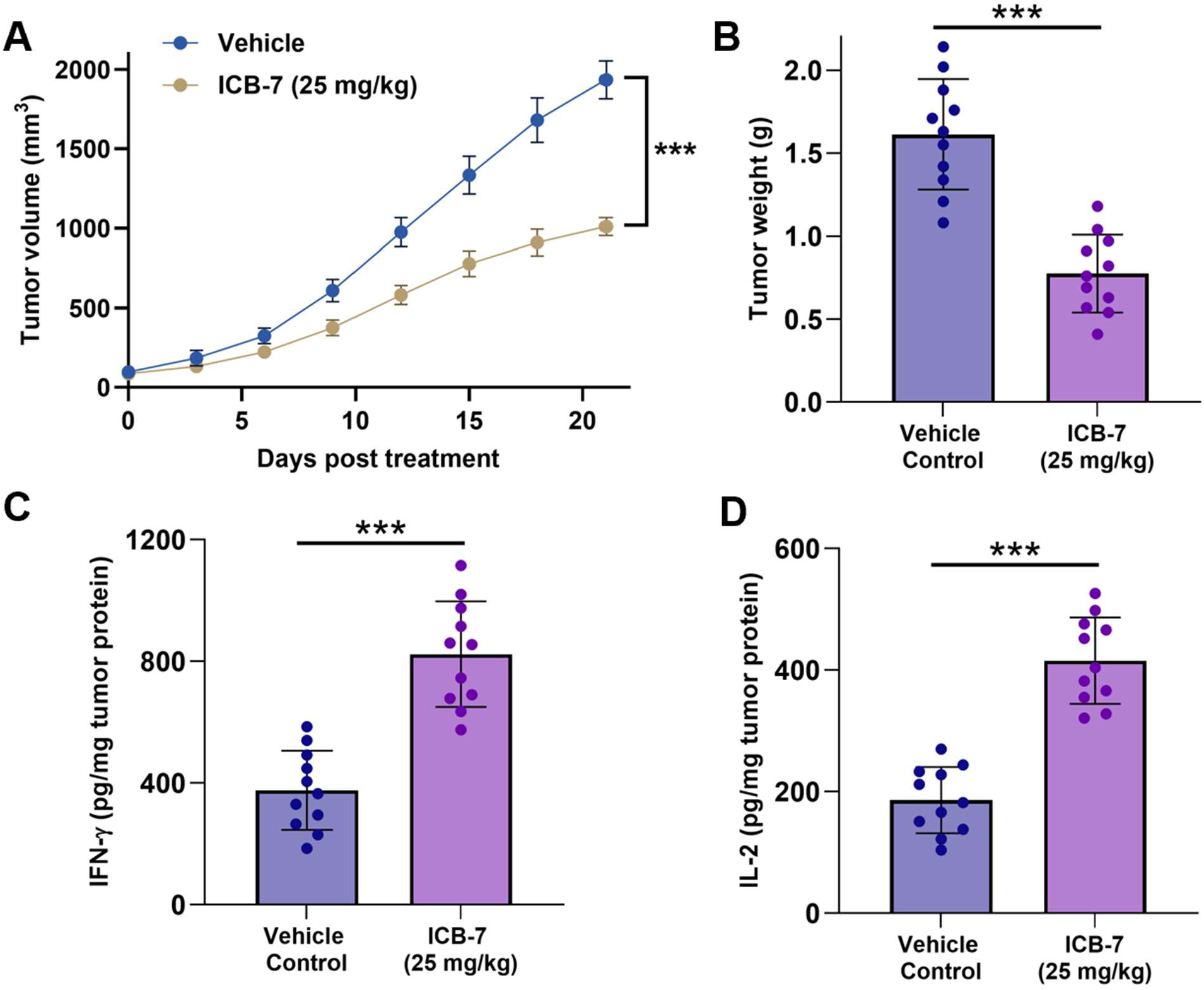
ICB-7 suppresses CT26 tumor growth and promotes anti-tumor immune activation in vivo. **(A)** Tumor growth curves of CT26 syngeneic colorectal tumors established in BALB/c mice treated with vehicle or **ICB-7** (25 mg/kg, PO, QD). Treatment was initiated once tumors reached approximately 75-100 mm^3^. **ICB-7** significantly inhibited tumor progression throughout the study period. **(B)** Final tumor weights measured at study termination demonstrating reduced tumor burden following **ICB-7** treatment. **(C,D)** ELISA analysis of tumor lysates revealed elevated intratumoral IFN-γ and IL-2 levels in tumors from **ICB-7**-treated mice relative to vehicle controls, consistent with enhanced immune activation within the tumor microenvironment. Data are presented as mean ± SEM with individual animals shown. Statistical significance was determined using an unpaired two-tailed Student’s t-test. \*\*\**p* < 0.001.

## CONCLUSION

In conclusion, this study demonstrates ILT3/LILRB4 as a tractable target for small molecule immunomodulation and reveals the feasibility of pharmacologically reprogramming suppressive myeloid immunity using direct ILT3 binders. Through a Dianthus/TRIC-based discovery workflow, we identified small molecules capable of directly engaging ILT3, disrupting the ILT3-SCG2 signaling axis, and reversing suppressive immune phenotypes across multiple functional systems. The lead compound, **ICB-7**, demonstrated robust biophysical target engagement, inhibition of downstream SHP1/SHP2/STAT3 signaling, restoration of anti-tumor immune activity in patient-derived co-culture models, and significant anti-tumor efficacy in vivo following oral administration. Computational studies further revealed a previously uncharacterized ligand-binding region within the ILT3 D2 domain, providing structural insight into small molecule recognition of this immune checkpoint receptor. Collectively, these findings provide proof-of-concept that suppressive myeloid checkpoints can be effectively targeted using small molecules and support further development of ILT3-directed immunomodulators as a promising therapeutic strategy for cancer immunotherapy.

## EXPERIMENTAL

### Compound Procurement

All compounds evaluated in this study were commercially available compounds purchased from Enamine (Monmouth Junction, NJ, USA). Enamine IDs for all validated hits from this study are provided in Table S1.

### Dianthus TRIC Assay

Initial screening of the Enamine Protein Mimetic Library (8,961 compounds) for ILT3 binding was performed using the temperature-related intensity change (TRIC) assay on a Dianthus NT.23 Pico instrument (NanoTemper Technologies). Recombinant human His-tagged ILT3 protein (from SinoBiological, Catalog# 16742-H08H) was fluorescently labeled with RED-tris-NTA 2nd Generation dye (NanoTemper, Cat. #MO-L018) according to the manufacturer’s protocol.

Labeled ILT3 (final concentration: 30 nM) was incubated with test compounds at a fixed concentration (10 µM) in assay buffer (PBS pH 7.4 supplemented with 0.05% Tween-20 and 1% (v/v) DMSO). Fluorescence was measured in Dianthus 384-well plates (NanoTemper Technologies, Catalog# DI-P001A) with LED power set to 40%. Fluorescence was recorded at 670 nm and 650 nm, and normalized fluorescence (F_norm_) was calculated as the ratio of F670/F650. Binding responses were quantified as Fnorm. Compounds with ΔF_norm_ ≥ 3 units and reproducible responses across replicates (n = 5) were classified as hits.

### Quantitative Binding Affinity Determination Using Monolith

Binding affinities of selected hits were quantified by microscale thermophoresis using a Monolith X instrument (NanoTemper Technologies). His-tagged ILT3 protein was labeled with RED-tris-NTA 2nd Generation dye using the Monolith His-Tag Labeling Kit (Cat. #MO-L018) following the manufacturer’s guidelines. Compound titrations were prepared as serial dilutions (PBS buffer, pH 7.4, 0.05% Tween-20, 1% (v/v) DMSO). Following a 15 min incubation at room temperature in the dark, samples were loaded into Monolith capillaries (Cat. #MO-K022) and analyzed at 25 °C using 40% LED power and medium MST power settings. Normalized fluorescence (F_norm_) values were determined as the ratio of fluorescence intensity after and before IR laser heating. Each compound was evaluated in five technical replicates. Dissociation constants (K_D_) were calculated from three independent experiments using MO.Affinity Analysis software and GraphPad Prism 10, applying standard dose-response fitting models. Data represent mean ± SD (n=5).

### TR-FRET ILT3-SCG2 Competition Assay

The ILT3-SCG2 interaction was measured using a homogeneous TR-FRET assay. Recombinant human ILT3-ECD-Fc and His-tagged SCG2 were incubated with terbium cryptate-labeled anti-His antibody (donor) and XL665-labeled anti-human Fc antibody (acceptor) in medium-binding white 384-well plates (Greiner, #784075). Recombinant human SCG2 carrying C-terminal polyhistidine tag was purchased from SinoBiological (catalog number 13441-H08H), while the ECD of human ILT3 fused to the Fc region of human IgG1 was obtained from SinoBiological (catalog number 16742-H02H). Terbium cryptate-labeled anti-His monoclonal antibody as the donor fluorophore and an XL665-conjugated anti-human Fc antibody as the acceptor, both sourced from Revvity (catalog numbers 61HISTLF and 61HFCXLF, respectively). Final assay concentrations were 20 nM ILT3-ECD-Fc, 20 nM SCG2-His, 1 nM donor antibody, and 10 nM acceptor antibody in PBS containing 0.1% DMSO. **ICB-7** was added as a serial dilution and plates were incubated for 2 h at room temperature.

TR-FRET signals were measured using a Tecan Spark plate reader (340 nm excitation; 620 nm donor emission; 665 nm acceptor emission). Data acquisition parameters included 50 flashes per well, a 50 µs delay time, and a 400 µs integration window. A blocking anti-ILT3 antibody (h128-3, catalog number HV126013) was used as a positive control. Percent inhibition values were normalized to DMSO-treated controls and fitted using nonlinear regression in GraphPad Prism to determine IC_50_ values.

**CETSA.** CHO cells stably expressing human ILT3/LILRB4 (from Creative Biogene, Cat# CSC-RO0667) were maintained in Ham’s F-12 medium supplemented with 10% fetal bovine serum and 1% penicillin-streptomycin under standard humidified culture conditions (37 °C, 5% CO2). Cells were seeded in 384-well plates at 12,000 cells/well and incubated overnight at 37 °C in 5% CO₂. **ICB-7** was added at the indicated concentrations and incubated for 60 min. Cells were subsequently subjected to thermal challenge at 51 °C for 3 min (using a calibrated Bio-Rad C1000 Touch thermal cycler), followed by cooling to room temperature. Following thermal challenge, cells were lysed directly in assay plates by addition of lysis buffer supplemented with protease inhibitor cocktail and incubated for 30 min at 4 °C with gentle agitation. Lysates were subsequently clarified by centrifugation at 4000 × g for 10 min to remove aggregated and insoluble protein species generated during thermal denaturation.

The soluble fraction of ILT3 remaining after thermal challenge was quantified using an AlphaLISA-based immunodetection format. Briefly, clarified lysates were incubated with ILT3-specific biotinylated detection antibody and AlphaLISA acceptor beads according to the manufacturer’s instructions, followed by addition of streptavidin-coated donor beads under reduced-light conditions. After incubation at room temperature, AlphaLISA signal was measured using Tecan Spark plate reader. CETSA stabilization signals were normalized to vehicle-treated controls, and dose-response curves were generated using nonlinear regression analysis. Data represent mean ± SD, n = 5.

### Computational study

Two compounds identified through virtual screening, namely **ICB-7** and **ICB-9**, were selected for molecular modeling investigations in order to elucidate their potential interaction mode(s) with the target protein. The three-dimensional structures of both compounds were generated using the Avogadro software package.^47^

The structure of Leukocyte Ig-like Receptor LILRB4 (ILT3) is available in the Protein Data Bank under the accession code 3P2T. This X-ray crystallographic structure^48^ was determined at a high resolution of 1.7 Å and does not contain any missing residues. The protein consists of a single polypeptide chain divided into two domains, D1 and D2, and contains three disulfide bridges.

To investigate the possible binding modes of **ICB-7** and **ICB-9**, onto ILT3 molecular docking calculations were performed using AutoDock 4.2.^49^ A cubic docking grid corresponding to a volume of 229.22 nm^3^ was defined around the ILT3 protein in order to explore all potential ligand-binding regions. To ensure statistically reliable docking results, 20 independent docking runs, each generating 100 binding poses, were performed, thereby allowing an extensive exploration of the ILT3 protein surface. The resulting docking structures were then clustered according to a RMSD lower than 2 Å. The conformation selected was the one which presented the lowest docking free energy of binding in the most populated cluster.^50^

All MD simulations were performed using the Amber software package.^51,52^ The GAFF2 force field^53^ was employed with AM1-BCC partial atomic charges^54,55^ along with the protein ff19SB force field.^56^ Water molecules were described using the TIP3P model,^57^ and all simulations were carried out under periodic boundary conditions. Each simulation started with 20,000 steps of geometry optimization, followed by 500 ns of molecular dynamics simulations performed under NPT conditions at 30 °C. Trajectory analyses were performed for the last 100 ns of each simulation, considering that equilibration is reached, using the cpptraj^58^ and VMD^59^ software. This last software was also used for images generation along with ChimeraX^60^. Free energy of interaction was computed with the MMGBSA technique^61^.

### THP-1-Derived Macrophage-Like Cell Signaling Assays

THP-1 cells (from ATCC) were seeded at 1 × 10^6^ cells/well in 6-well plates and differentiated into macrophage-like cells by treatment with phorbol 12-myristate 13-acetate (PMA, 100 nM) for 48 h, followed by a 24 h resting period in PMA-free complete RPMI-1640 medium supplemented with 10% fetal bovine serum and 1% penicillin–streptomycin. Differentiated cells were stimulated with recombinant human SCG2 (100 ng/mL) in the presence of increasing concentrations of **ICB-7** for 1 h at 37 °C. Cells were subsequently washed with ice-cold PBS and lysed using RIPA buffer supplemented with protease and phosphatase inhibitor cocktails. Lysates were clarified by centrifugation at 14,000 × g for 15 min at 4 °C prior to downstream phospho-signaling analysis.

Phosphorylated SHP1, phosphorylated SHP2, and phosphorylated STAT3 were quantified using commercial phosphospecific ELISA kits (abcam, catalog# ab279924, ab314344, and ab279941, respectively) according to the manufacturers’ protocols. Lysates were normalized for total protein content using a BCA assay before loading. Data were normalized to SCG2-treated controls and expressed as percent of SCG2-induced signaling. IC_50_ values were calculated using nonlinear regression. Data represent mean ± SD, n = 5.

### Human Co-Culture Assays

For colorectal cancer co-culture assays, HCT116 or HT-29 cells (from ATCC) were seeded in flat-bottom 96-well plates at 1 × 10^4^ cells/well and allowed to adhere overnight in complete RPMI-1640 medium supplemented with 10% fetal bovine serum and 1% penicillin–streptomycin. PBMCs isolated from colorectal cancer patients (from STEMCELL Technologies) by Ficoll density-gradient centrifugation were added at an effector-to-target ratio of 10:1 in the presence or absence of recombinant human SCG2 (100 ng/mL) and increasing concentrations of **ICB-7** (1-25 μM). A blocking anti-LILRB4 antibody (h128-3, catalog number HV126013) was included as a target-specific positive control. Co-cultures were incubated for 72 h at 37 °C in 5% CO_2_. Supernatants were collected for IFN-γ and IL-2 quantification by ELISA (abcam catalog# ab46025 and #ab270883, respectively), and tumor-cell viability was assessed using CellTiter-Glo (from Promega) according to the manufacturer’s protocol.

For AML co-culture assays, THP-1 cells (from ATCC) or primary AML blasts (from STEMCELL Technologies) were co-cultured with PBMCs from AML patients (from STEMCELL Technologies) or enriched CD3^+^ T cells under analogous conditions. Supernatants were collected for IFN-γ and IL-2 ELISA analysis (abcam catalog# ab46025 and #ab100706, respectively). Cytotoxic T-cell activation was assessed by measuring Granzyme B^+^ CD8^+^ T cells using flow cytometry. Tumor-cell viability was quantified using CellTiter-Glo (from Promega). Data were normalized to media or SCG2-treated controls as indicated in the figure legends. Data represent mean ± SD, n = 5.

### PK studies

In vitro PK profiling was performed as we previously reported.^40,41^ In vivo PK studies were conducted in female BALB/c mice (6-8 weeks old; n = 5 per time point). The tested compound was administered by intravenous injection (2 mg/kg) or oral gavage (20 mg/kg) in a formulation consisting of 5% DMSO, 40% PEG400, and 55% sterile saline (v/v/v), prepared fresh prior to dosing. Blood samples were collected at 0.25, 0.5, 1, 2, 4, 8, and 24 h post-dose via tail vein sampling into EDTA-coated tubes. Plasma was isolated by centrifugation at 3,000 × g for 10 min at 4 °C.

### CT26 Syngeneic Colorectal Tumor Model

All animal studies were performed in accordance with protocols approved by the Institutional Animal Care and Use Committee (IACUC). Female BALB/c mice (6-8 weeks old) were obtained from Jackson Laboratory and acclimated for at least 1 week prior to experimentation under standard pathogen-free housing conditions with ad libitum access to food and water.

CT26 murine colorectal carcinoma cells were maintained in RPMI-1640 medium supplemented with 10% fetal bovine serum and 1% penicillin-streptomycin. Cells were harvested during logarithmic growth phase, washed twice with sterile PBS, and resuspended in cold PBS. Each mouse received a subcutaneous injection of 5 × 10^5^ CT26 cells into the right flank in a final injection volume of 100 µL. Tumor growth was monitored three times weekly using digital calipers. Once tumors reached approximately 75-100 mm^3^, mice were randomized into treatment groups (n = 11 mice/group) to ensure comparable mean starting tumor volumes across cohorts. **ICB-7** was formulated fresh daily in vehicle consisting of 5% DMSO, 40% PEG400, and 55% sterile saline (v/v/v). Mice received vehicle control or **ICB-7** at 25 mg/kg by oral gavage once daily for 21 consecutive days. The selected dosing regimen was guided by in vivo PK studies demonstrating sustained plasma exposure above the biochemical and cellular potency range associated with ILT3 signaling inhibition.

Body weight and clinical condition were monitored throughout the study to assess tolerability. Humane endpoints included tumor ulceration, impaired mobility, or tumor burden exceeding institutional limits. At study termination, mice were euthanized by CO_2_ inhalation followed by cervical dislocation. Tumors were excised, weighed, photographed, and snap-frozen for downstream pharmacodynamic analyses.

Tumor lysates were prepared using ice-cold lysis buffer supplemented with protease inhibitors and clarified by centrifugation. Total protein concentrations were quantified using a BCA assay prior to normalization. Intratumoral IFN-γ and IL-2 levels were quantified using commercial ELISA kits (abcam catalog# 100689 and #ab46096, respectively) according to the manufacturers’ protocols. Cytokine levels were normalized to total tumor protein content and expressed as pg/mg tumor protein.

### Statistical Analysis

Data are presented as mean ± SD or mean ± SEM as indicated in the figure legends. Statistical analyses were performed using GraphPad Prism. For two-group comparisons, unpaired two-tailed Student’s t-tests were used. For multiple-group comparisons, one-way ANOVA followed by Tukey’s post hoc test was applied. Tumor growth curves were analyzed using repeated-measures ANOVA where appropriate. *P* values <0.05 were considered statistically significant.

