## Supporting Information for "Targeting the Myeloid Immune Checkpoint ILT3 (LILRB4) with Small Molecules Enables Reprogramming of Suppressive Tumor Immunity"

| **Contents** | |  |
| --- | --- | --- |
| Chemical structures of the validated hit compounds identified by Dianthus screening | | S2 |
| **ICB-7** restores anti-tumor immune activity across colorectal cancer and AML co-culture systems | | S3 |
| **ICB-7** directly binds recombinant murine ILT3 as determined by MST  Selectivity profiling of **ICB-7** across immune checkpoint and LILR family proteins | | S4  S5 |
| HRMS chart for **ICB-7** | | S6 |
| HPLC purity trace for **ICB-7** | | S6 |

**Table S1**. Chemical structures of the validated hit compounds identified by Dianthus screening.

| **Compound name** | **Chemical structure** | **Enamine ID** | **LILRB4 (K_D_)** |
| --- | --- | --- | --- |
| **ICB-7** | 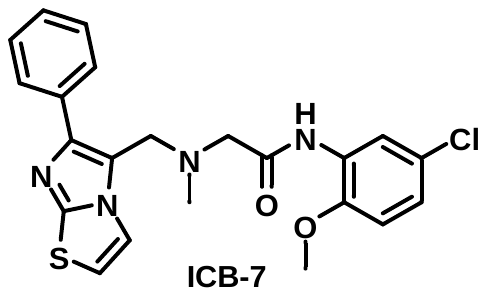 | **Z89028154** | 156 ± 11.8 nM |
| **ICB-9** | 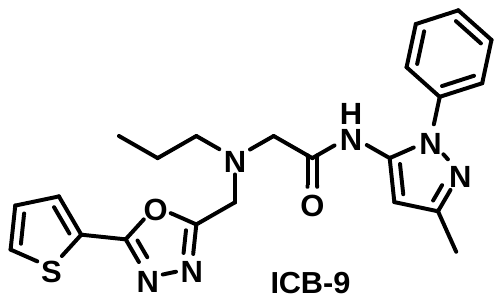 | **Z89143671** | 887 ± 36.1 nM |
| **ICB-10** | 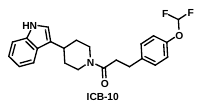 | **Z317731376** | 12.7 ± 1.41 μM |
| **ICB-12** | 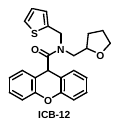 | **Z91156675** | 25.2 ± 5.72 μM |

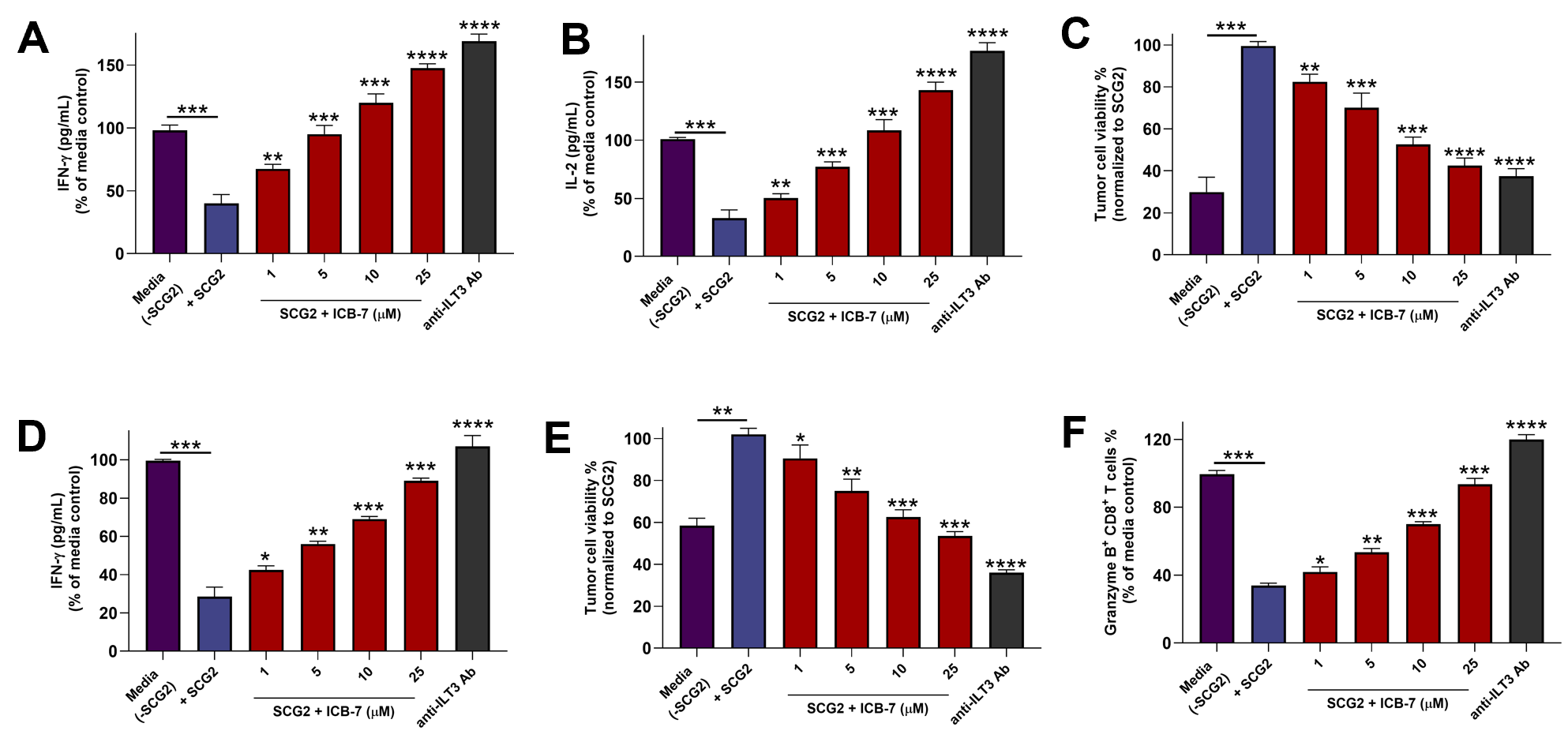

**Figure S1. ICB-7 restores anti-tumor immune activity across colorectal cancer and AML co-culture systems. (A,B)** In colorectal HT-29 cancer co-cultures, SCG2 stimulation suppressed anti-tumor immune activity, resulting in reduced IFN-γ and IL-2 secretion. Treatment with **ICB-7** restored cytokine production in a concentration-dependent manner. **(C) ICB-7** significantly reduced colorectal HT-29 cancer cell viability in SCG2-treated co-cultures, consistent with restoration of immune-mediated tumor killing. **(D)** In THP-1 AML co-cultures, **ICB-7** restored IFN-γ secretion suppressed by SCG2-mediated ILT3 signaling. **(E) ICB-7**  increased the frequency of Granzyme B^+^ CD8^+^ T cells in a dose-dependent manner, indicating recovery of cytotoxic T-cell function. **(F) ICB-7** significantly reduced THP-1 AML viability in SCG2-treated co-cultures. Data represent mean ± SD from independent donors performed in technical replicates. Statistical significance was determined using one-way ANOVA with Tukey’s multiple comparisons test. **p* < 0.01, ***p* < 0.05, ****p* < 0.001, *****p* < 0.0001 relative to media +SCG2.

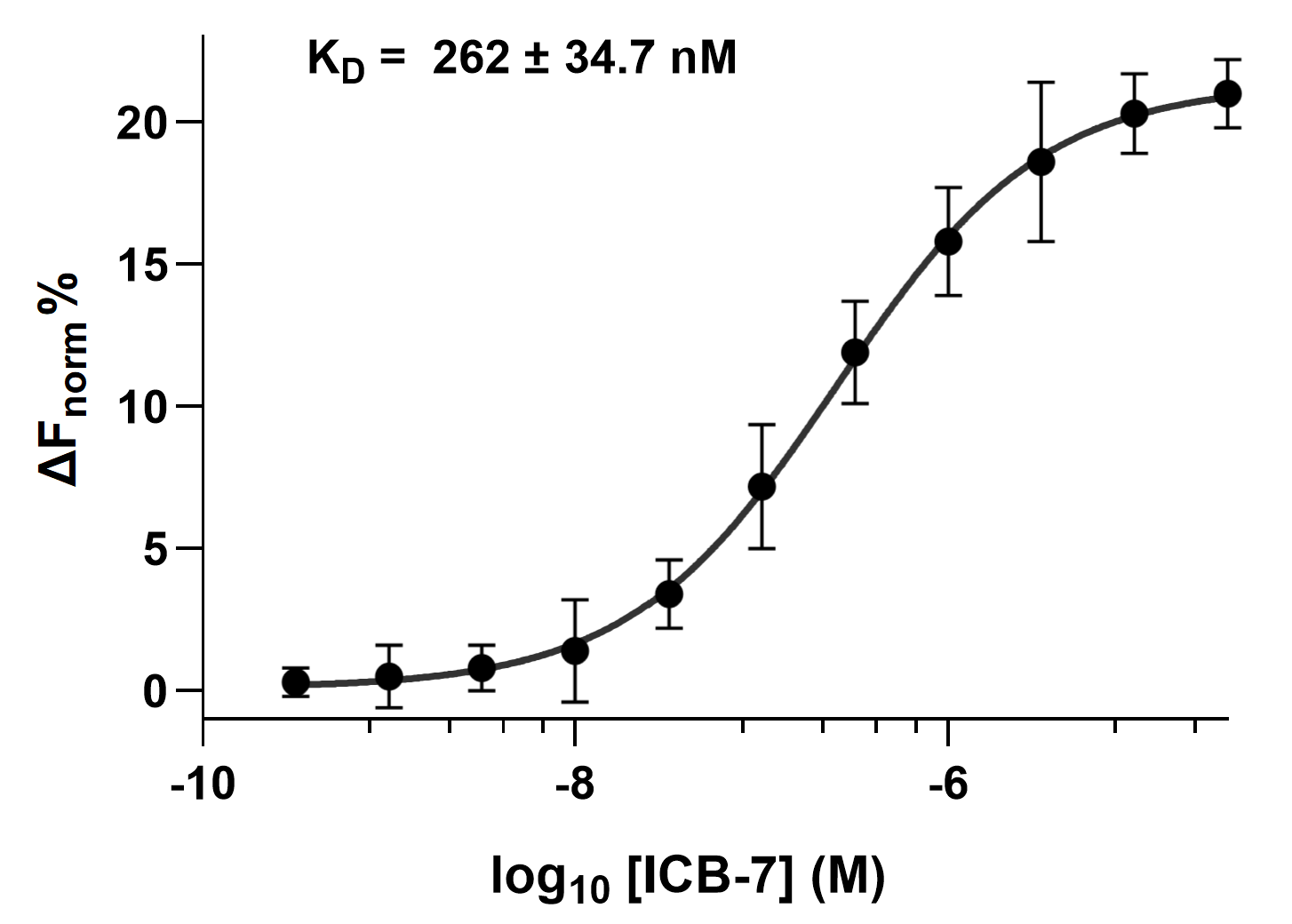

**Figure S2. ICB-7 directly binds recombinant murine ILT3 as determined by MST.** Binding of **ICB-7** to recombinant murine ILT3 was evaluated using MST under solution-phase conditions. **ICB-7** exhibited concentration-dependent interaction with murine ILT3. Data represent mean ± SD (n=5).

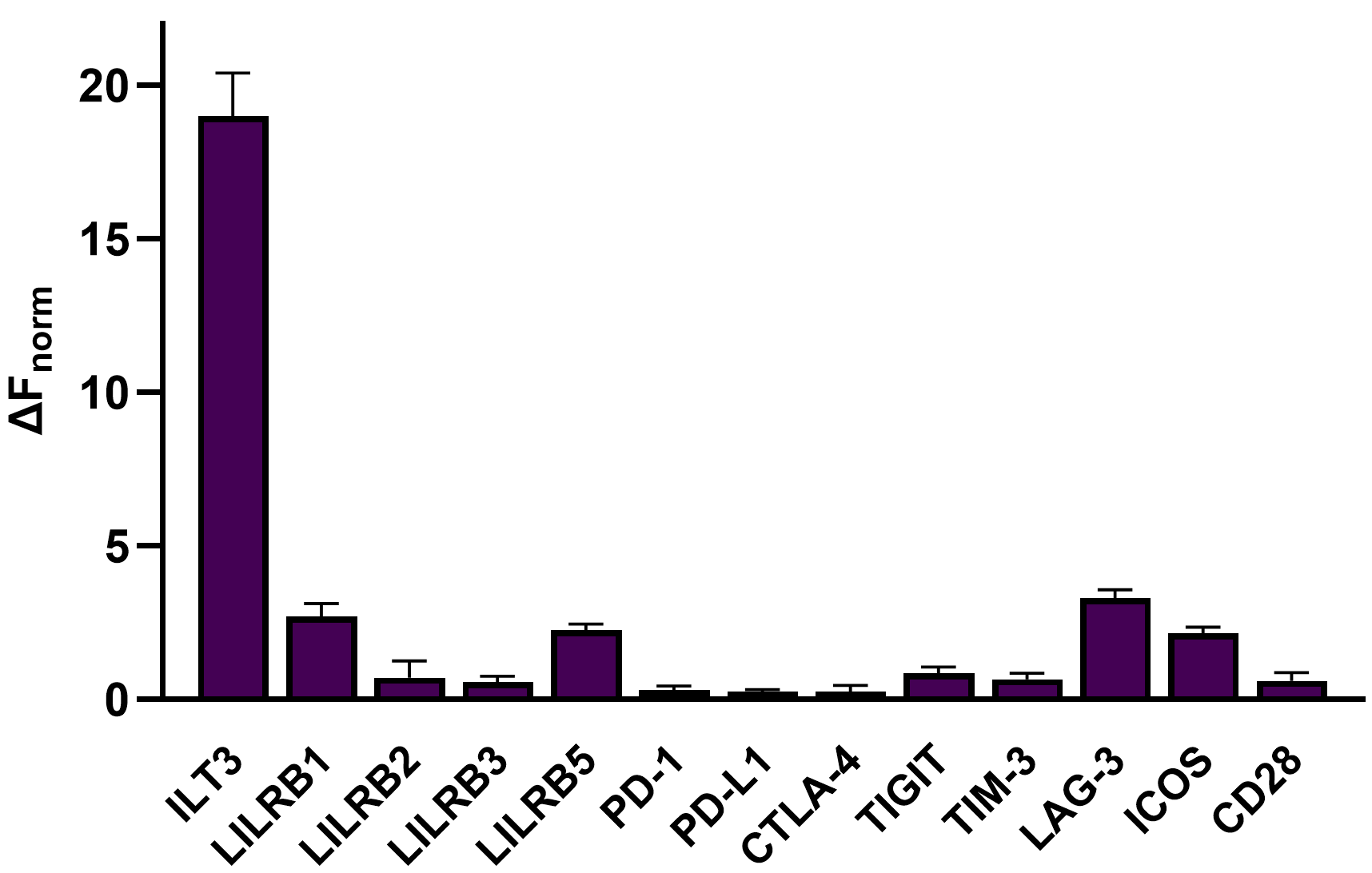

**Figure S3. Selectivity profiling of ICB-7 across immune checkpoint and LILR family proteins.**
Dianthus/TRIC analysis showing normalized fluorescence changes (ΔF_norm_) following incubation of **ICB-7** (10 μM) with a panel of immune checkpoint proteins and related LILR family members. Data represent mean ± SD (n = 5).

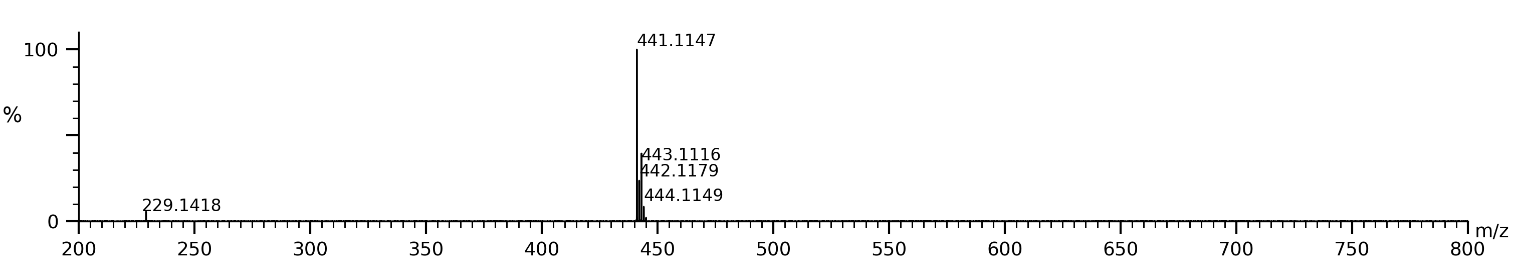

**Figure S4. HRMS chart for ICB-7.**

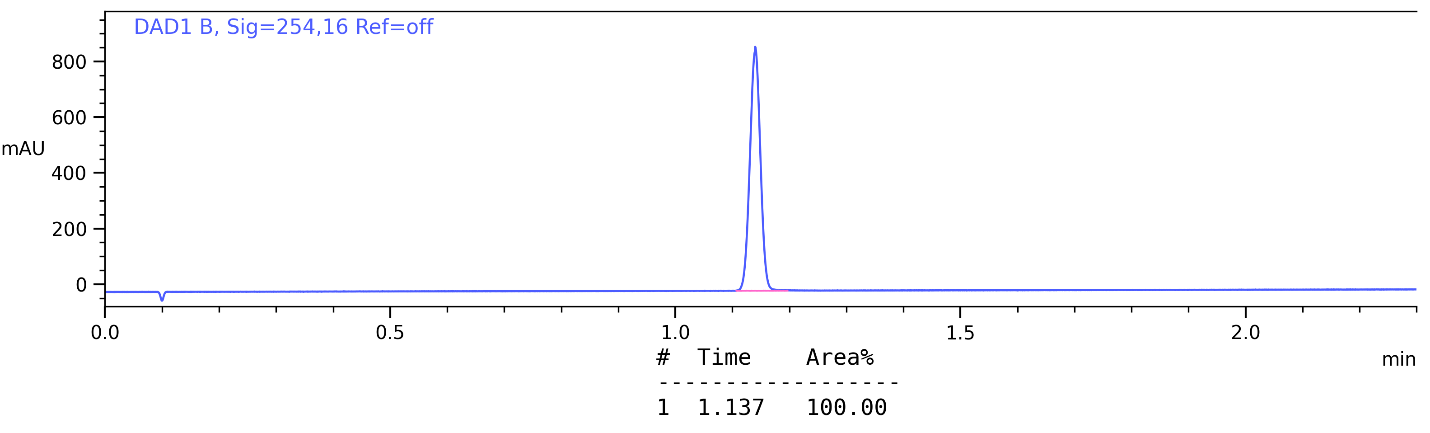

**Figure S5. HPLC purity trace for ICB-7.**
